# Capturing sleep-wake cycles by using day-to-day smartphone touchscreen interactions

**DOI:** 10.1101/479014

**Authors:** Jay N. Borger, Reto Huber, Arko Ghosh

## Abstract

Body movements drop with sleep and this behavioural signature is widely exploited to infer sleep duration. However, a reduction in body movements may also occur in periods of intense cognitive activity and the ubiquitous use of smartphones may capture these wakeful periods otherwise hidden in the standard measures of sleep. Here we continuously captured the gross body movements using standard wrist-worn accelerometers to quantify sleep (actigraphy) and logged the timing of the day-to-day touchscreen events (‘tappigraphy’). Using these measures, we addressed how the gross body movements overlap with the cognitively engaging digital behaviour (from n = 84 individuals, accumulating 1384 nights). We find that smartphone use was distributed across a broad spectrum of physical activity levels but consistently peaked at rest. We estimated the putative sleep onset and wake-up times from the actigraphy data to find that these times were well correlated to the estimates from tappigraphy (R^2^ = 0.9 for sleep onset and wake-up time). However, actigraphy overestimated sleep as virtually all of the users used their phones during the putative sleep period. Interestingly, the probability of touches remained greater than zero for ~ 2 h after the putative sleep onset and ~ 2 h before the putative wake-up time. Our findings suggest that touchscreen interactions are widely integrated into modern sleeping habits – surrounding both sleep onset and waking-up periods – yielding a new approach to measuring sleep. Smartphone taps can be leveraged to update the behavioural signatures of sleep with these peculiarities of modern digital behaviour.

## Introduction

There is a well-recognized need for tracking sleep in patients as well as in the general population. This rush to quantify sleep is partly driven by the increased awareness that sleep is crucial for cognitive performance and wellbeing. Body movements offer an easy proxy for sleep measurements. Essentially, users wear an accelerometer at the wrist or ankle and the recorded accelerations i.e., body movements are automatically converted into estimates of sleep ^1,2^. This popular method relies on the sharp drop in motility at sleep onset and the sharp rise with wakefulness ^1,3^. The body movements recorded at the joints may involve different levels of cognitive engagement – from reflexes to the postural control accompanying fine motor control. However, the extent of cognitive engagement does not enjoy a simple linear relationship with the movements recorded at the joints. For instance, the now common fine finger movements on the touchscreen are cognitively engaging but they may result in no or negligible signal deflections at the wrist. This lack of a simple relationship and the widespread use of smartphones in modern behaviour warrant an up-to-date perspective on tracking sleep based on motor activity.

In general, modern digital interactions offer unprecedented opportunities to quantify behaviour in the real world with major repercussions for sleep. For instance, the timing of social messaging such as on Twitter can be used to elaborate the diurnal behavioural patterns ^4,5^. This measure of online activity is somewhat limited in terms of capturing the behaviour continuously. For instance, only a fraction of the digital behaviour occurs via the social messaging server. Another approach has focused on the mobile device itself, and sleep-wake cycles can be inferred by machine learning algorithms that use the hardware state of the smartphone (i.e., phone on the charger and the screen being on or off) as inputs and sleep diaries as the ground truth ^6^. This ‘black-box’ approach is not designed to improve the fundamental understanding of motor behaviour and sleep, but it does underscore the value of capturing data from the smartphone sensors in the context of sleep. However, there is a large gap between the accuracy of phone-based sleep detection and the objective measures of sleep ^7^. Regardless of the current limitations and pending validations of these novel approaches, they do promise an economical, easy to administer and highly scalable measure of sleep based on existing sensors in contrast to approaches that require extra sensors as used in standard actigraphy.

In this study, we used standard wrist-worn actigraphy to quantify sleep-wake cycles and in parallel recorded the time-stamps of smartphone touchscreen interactions. Although smartphones have built-in accelerometers capable of monitoring body movements – as long as it is carried by the user – the wrist-worn approach ensures that all of the movements are independently and continuously recorded including when at asleep. We focused on the Cole-Kripke algorithm which is well studied and widely used to infer sleep from the body movements. This algorithm has been validated against the gold-standard or direct measure of sleep using polysomnography^3,8–10^. By merging these distinct measures, we quantify the patterns of overlap between smartphone behaviour and sleep and validate a novel approach to measure sleep derived from the smartphone interactions alone.

## Methods

### Participants and recruitment

Participants across the campus of Leiden University were recruited by using advertisements on a closed online platform and department-wide emails. Candidates with known neurological and psychiatric diagnosis based on self-reports were blocked from recruitment. Due to technical limitations, those users with an Android operating smartphone were invited to participate and under the condition that the phone remains strictly un-shared during the study period. A total of 84 right-handed participants were recruited (44 females, 16-45 years of age, mean age 23). The experimental procedures used here were approved by the Ethical Committee at the Institute of Psychology at Leiden University. All the participants provided written and informed consent and were compensated for their time using a cash reward or course credits. The weight with one layer of clothing, height and the year/month of birth was collected from each participant.

### Actigraphy measurement

The gross movements (3-axis accelerometer), the ambient light and near body temperature were measured using GENEACTIV watches (Activinsights, Cambridgeshire, UK). Participants were instructed to wear the watch on both wrists for a minimum of 2 weeks, and the data from the left wrist is primarily presented here. One individual discontinued wearing the watches after a period of 5 days due to a skin rash, another 3 individuals discontinued after 4 days due to difficulties in falling asleep wearing the watches and another individual discontinued after 2 days due to employer restrictions on watches in general. The watches were set to acquire the data at 50 Hz and the data was recovered after 14 days of acquisition, only to be reset for continued use if the subjects were willing to participate for an additional week.

### Tappigraphy measurement and on-phone sleep diary

The touchscreen interactions were quantified using the TapCounter App (QuantActions Ltd. Lausanne, Switzerland). The App was installed by each user from the Google Playstore (Google, Mountain View, USA). The App is designed to gather the precise timestamps of all touchscreen interactions and operates in the background. Only those touchscreen interactions which occurred during the ‘unlocked’ state of the screen were considered here. Each user was provided with a unique user code – and when entered into the App the data was streamed to the cloud along with the unique code for further processing. All data was encrypted during transmissions. Users were instructed to note the bed, sleep, wake-up and out-of-bed times every day during the actigraphy measurements on a ‘notes’ feature built into the TapCounter.

### Actigraphy algorithm

The accelerations gathered along the three axes by the actigraphy watches were combined using the sum of squares and low-pass filtered at 2 Hz. To estimate the putative sleep and wake times, we employed the standard Cole-Kripke algorithm on the filtered data with slight modifications^3^. A key part of this algorithm – the minute-by-minute categorization of the data into rest-active states based on the weighted sum of the current minute with that of the surrounding minutes – was extracted to study the physical activity state during smartphone usage. The algorithm was implemented on MATLAB (MathWorks, Natick, USA) and used pre-existing codes^11^. Essentially through the modifications, the automatic scoring of sleep and wake by the Cole-Kripke algorithm was further checked by the near body temperature and ambient light measurements. Firstly, any putative sleep period where the median temperature dropped below 25^ο^C was ignored – removing instances where the user removed the watch from the body. Secondly, any putative sleep period where the median ambient light levels failed to drop below 25 lux was ignored and thus restricting the analysis to nighttime sleep and ignoring day-time naps. Thirdly, the putative sleep times had to contain a 10% (36 min) overlap with the 6 h low-activity period determined using a 24h sinewave fit (Casey Cox’s *cosinor* function implemented in MATLAB)^12^. This final step is commonly substituted using sleep diaries, but our approach avoided mixing the subjective diary entries with the objective measurements to determine sleep durations – ensuring the estimated durations are entirely objective.

### Tappigraphy algorithm

The touchscreen timestamps were processed using a separate algorithm designed to gather the gaps in smartphone use at the circadian rest phase (i.e., at the putative night). First, the phone data was reduced to binary states in 60s bins (1 as active and 0 as rest). The activity was further processed using a cut-off (5% in an hour threshold) such that the brief periods of activity surrounded by in-activity were labelled as rest. Next, we extracted all of the continuous gaps in smartphone activity, such that the gap in usage was greater than an arbitrarily set 2 h threshold. In a parallel set of computations, we obtained a 24 hour sinewave fit on the time-series of smartphone interactions using the Cosinor analysis (Casey Cox’s *cosinor* function implemented in MATLAB)^12^. This sinewave fit was then used to determine the 6-hour long periods with the least activity in the tapping data in 24-hour windows. The two parallel streams were combined to select those activity gaps which had a minimum of arbitrarily set 10% overlap (36 min) with the 6-hour period, and these gaps were labelled as ‘sleep’.

### Statistical analysis

Robust linear regressions (using the bi-square fitting method) was used to test relationships between tappigraphy and actigraphy, also to explore the relationship between tappigraphy ‘measurement errors’ and smartphone usage (implemented using the *fitlm* function in MATLAB). To enable the correlations of values from a 24h clock in a linear space a simple transformation for the sleep-onset values under 10 am, such that 01h in past midnight was considered as 25h. In t-tests, the *α* value was set 0.05 and was adjusted using Bonferroni correction. In the analysis of how the collected demographic information (age, gender, height and weight) reflect on sleep, the 2 subjects with ages higher than 35 were excluded as outliers (> 5STD from the mean).

## Results

### Physical activity and smartphone usage

We estimated the variations in smartphone interactions at the different levels of physical activity (Fig. 1). In keeping with our goal of better understanding sleep-wake cycles, we quantified the activity in terms of the actigraphy ‘D’ values, where D is proportional to the sum of the acceleration at any given minute and the surrounding minutes. Importantly, D < 1 corresponds to sleep in the Cole-Kripke algorithm^3^. In all of the participants, the probability of observing smartphone interactions increased when at physical rest (D ≈ 1, Fig. 1). In 42/87 participants, the smartphone interactions were maximum between the D values of 0-2 and 34/87 participants show the maximum values at greater than 6. Interestingly the smartphone behaviour at physical rest was related to the behaviour when physically active, such that the higher the smartphone usage at complete physical rest (0.25 ≥ D ≥ 0) the higher the phone usage when physically highly active (D > 7, R^2^ = 0.14, *β* = 1.13, *t*(72) = 3.33, *p* = 0.001).

**Figure 1.**
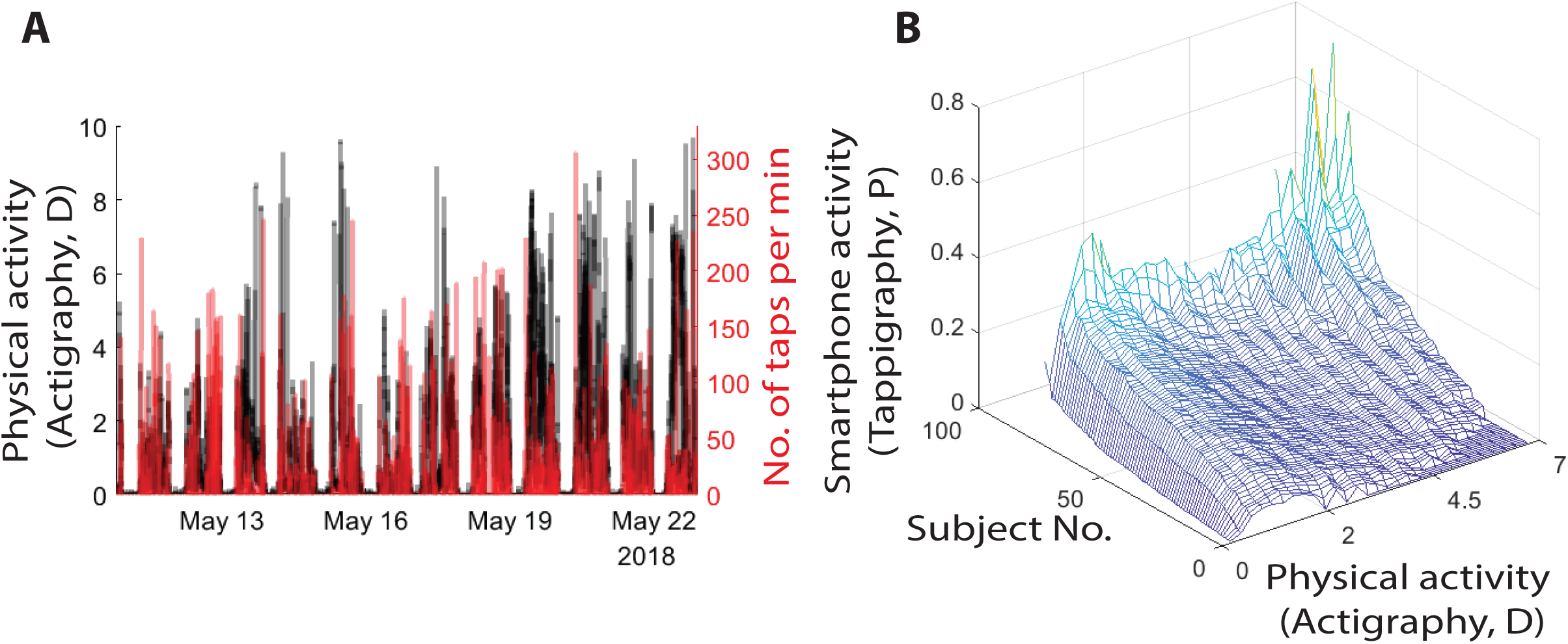
The relationship between smartphone touchscreen interactions and gross movements. (A) Data from a single participant showing the extent of the overlap between smartphone interactions quantified using an App running in the background (‘tappigraphy’) and overall physical activity measured at the wrist and quantified using an actigraphy algorithm, where D ≤ 1 is indicative of physical rest. (B) The probability of smartphone interactions in one-minute bins at different levels of physical activity quantified in steps of 0.25 D. The subjects are sorted according to the amount of smartphone activity at D = [0-0.25].

### Comparison of tappigraphy based sleep-wake estimations to standard actigraphy and sleep diaries

The high probability of smartphone touches at physical rest raised the opportunity that the putative sleep times can be simply estimated by observing the gaps in smartphone usage. We compared two standard methods used to estimate sleep with that of a new tappigraphy-based algorithm based on the gaps in smartphone usage. Pooling all of the measurements together, we found strong correlations between the putative sleep times determined by actigraphy vs. tappigraphy (Fig. 2). For the sleep onset times a linear fit with a slope ≈ 1 well captured the relationship (R^2^ = 0.85, *β* = 0.94, *t*(1382) = 86.85, *p* = 0). For the wake-up times there was strong correlation between the actigraphy (*x*) vs. tappigraphy (*y*) estimates as well (R^2^ = 0.9, *β* = 0.91, *t*(1382) = 112.02, *p* = 0). In terms of sleep duration, tappigraphy consistently ‘under-estimated’ sleep given that the slope was substantially below 1 (R^2^ = 0.29, *β* = 0.49, *t*(1325) = 22.9, *p* = 1 × 10^−98^). This regression yielded an intercept of 3.9 (*p* = 2.0 × 10^−85^) and *x* == *y* was at 7.6h, suggesting that the tappigraphy sleep durations > 7.6h are underestimates of sleep. This could not be simply explained by the fact that the data used was from the left wrist whereas the phones may be handled by the right. Using the right wrist movements, we again found a biased estimation of sleep duration (R^2^ = 0.32, *β* = 0.52, *t*(1375) = 23.7, *p* = 3.8 × 10^−104^ with an intercept of 3.8, *p* = 1.6 × 10^−77^). A similar pattern was found when comparing tappigraphy to sleep diaries. For sleep onset and wake-up times a linear fit with a slope ≈ 1 well captured the relationship between the two approaches (for sleep onset: R^2^ = 0.89, *β* = 0.10, *t*(1034) = 92.0, *p* = 0 and for wake-up times: R^2^ = 0.94, *β* = 0.99, *t*(1108) = 135.9, *p* = 0). As in actigraphy, the sleep duration was underestimated by tappigraphy (*y*) vs. diary (*x*, R^2^ = 0.59, *β* = 0.88, *t*(1023) = 38.5, *p* = 4.71 × 10^−201^) and the regression model had an intercept of 1.39 (*p* = 2.36 × 10^−14^). It is interesting to note how the sleep diary (*y*) related to actigraphy (*x*). The regression model was captured with a slope ≈ 0.5, suggesting subjects reported shorter durations compared to what was determined by actigraphy (R^2^ = 0.36, *β* = 0.47, *t*(1091) = 24.6, *p* = 7.42 × 10^−107^).

**Figure 2.**
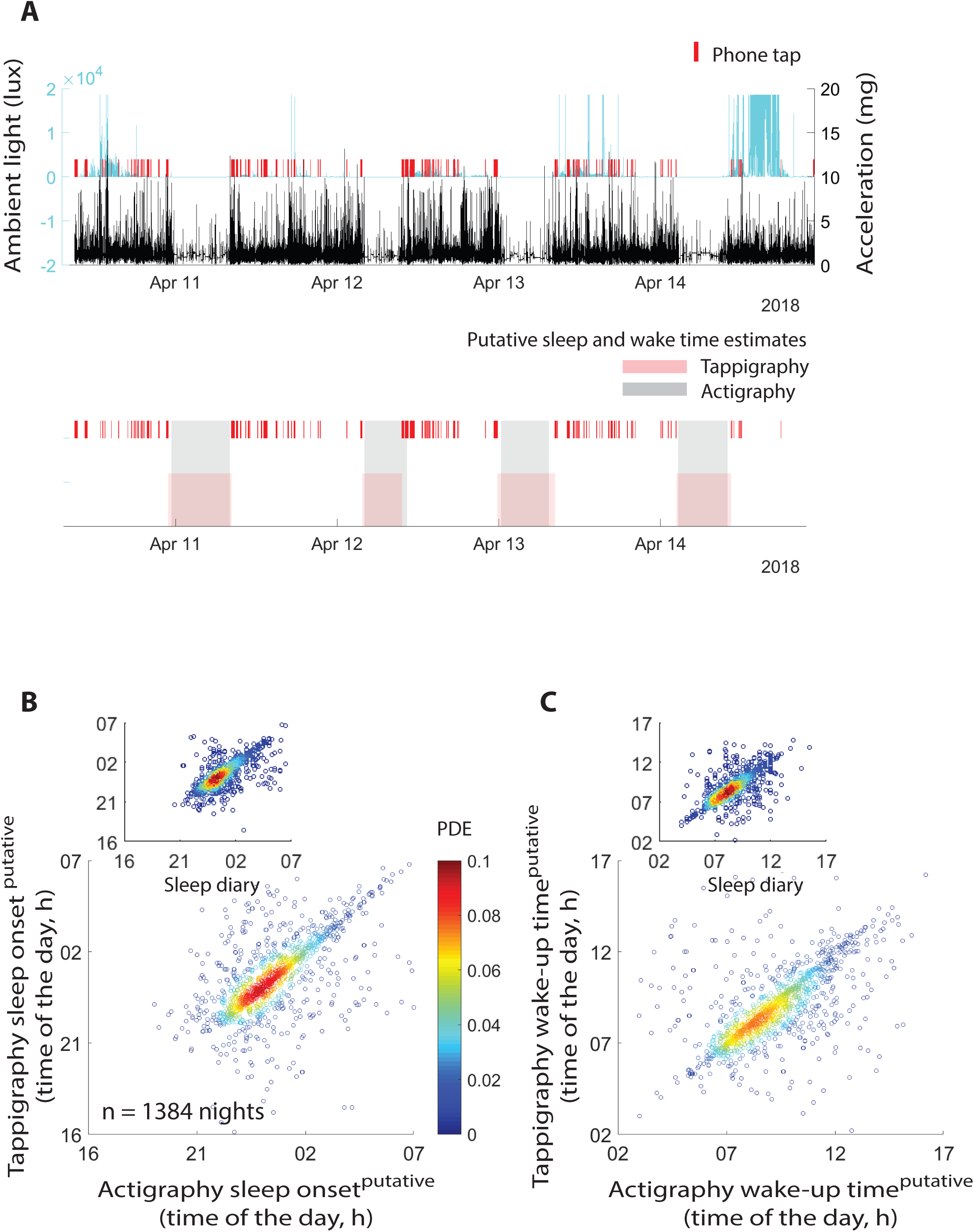
Comparison of tappigraphy based sleep estimates with that of actigraphy. (A) Actigraphy watches were used to quantify the amount of ambient light, near body temperature (not shown) and the body movements. The smartphone touches were simultaneously recorded by using an App running in the background. We used the Cole-Krpike algorithm to extract the putative sleep times from actigraphy and a new algorithm was designed to extract the sleep times from the smartphone touches (‘tappigraphy’). The relationship between the putative sleep onset times (B) and wake-up times (C) determined by using actigraphy vs. tappigraphy. Data was pooled across all the subjects. The time of the day is in local time and the inserts show the relationship between tappigraphy vs. sleep diaries.

### Inter-individual differences in actigraphy and tappigraphy-based sleep-wake estimations

A key question is how engaged must any user be on the smartphone for the tappigraphy based metrics to accurately reflect sleep. Considering the actigraphy based measures as ground truth, we found that the median sleep onset and wake-up times derived from tappigraphy were well correlated to the values obtained from actigraphy (Fig. 3, for sleep onset:, R^2^ = 0.72, *β* = 0.97, *t*(77) = 14.1, *p* = 4.45 × 10^−23^ and for wake-up times:, R^2^ = 0.60, *β* = 0.83, *t*(77) = 10.7, *p* = 6.00 × 10^−17^). Next, we determined the median percentage error in estimating sleep duration against the actigraphy values to find the median absolute % error to be 7.2, and median % error at -2.8, i.e., in the majority of the sampled population sleep was ‘under-estimated’ (negative error) by tappigraphy. Note that these negative errors may well mean that actigraphy over-estimates true sleep durations rather than tappigraphy underestimates the duration. This was further confirmed when comparing the population means derived by using actigraphy (8.5h ±0.94STD) vs. tappigraphy (8.1h ±1.1STD, *t*(78) = 2.4, *p* = 0.02). Interestingly, the errors were strongly related to smartphone usage – such that tappigraphy over-estimated sleep in comparison to actigraphy for users who generated a low number of touchscreen touches per day and the errors were reversed for the high smartphone users (R^2^ = 0.27, *β* = -0.0023, *t*(77) = - 5.23, *p* = 1.4 × 10^−6^). With the 0-error intercept set at ≈ 3200 touches per day. This value offers a guideline on the extent of engagement needed to estimate sleep using tappigraphy. As the number of available days of measurement varied from 5 to 31 days per individual, we opportunistically addressed whether the errors were linked to the number of days of measurement and this was not found to be the case (R^2^ = 0.02, *β* = -0.12, *t*(77) = - 0.62, *p* = 0.53).

**Figure 3.**
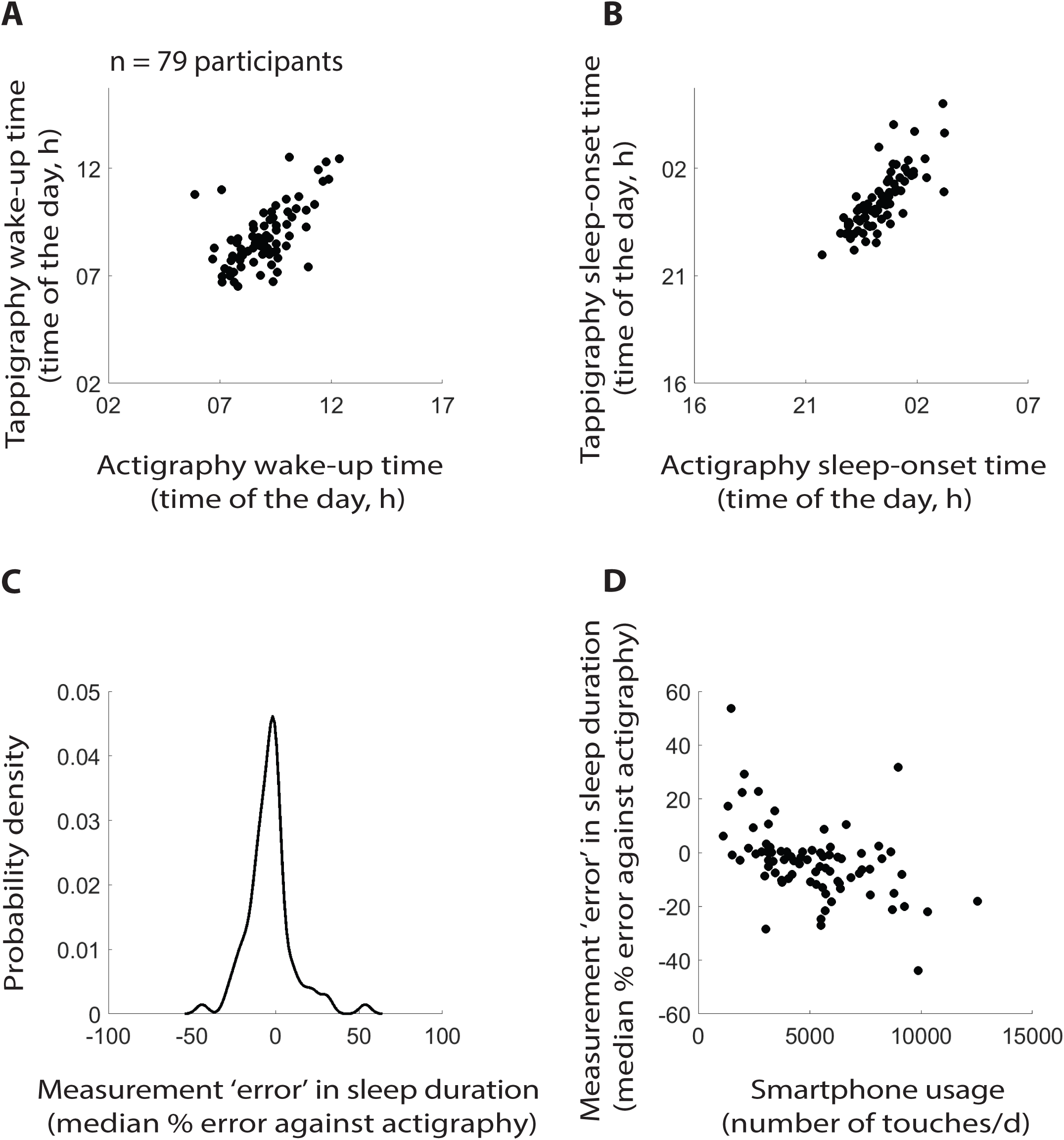
Population level comparison between tappigraphy and actigraphy. The central tendencies (median) of actigraphy base putative wake-up (A) and sleep (B) times were well correlated to the putative times determined using tappigraphy. (C) The population distribution of median measurement error in tappigraphy considering actigraphy as ground truth. (D) The relationship between the measurement error and smartphone usage.

As sleep may vary from one night to the next, we measured the intra-individual variation in sleep duration (CoV) to find that the actigraphy vs. tappigraphy values corresponded well to each other (R^2^ = 0.35, *β* = 0.57, *t*(77) = 6.13, *p* = 3.6 × 10^−8^). Finally, we exploited the demographic information to address whether the inter-individual differences in sleep durations and sleep CoV could be explained by the amount of phone usage (measured as number of touches per day), age, gender (dummy variable), height and weight. Age, height and weight were normally distributed, but the sample was focused on a rather narrow age-range (mean Age was 23 ±2.6 STD). For the actigraphy based sleep duration, the overall multiple regression model was not significant (R^2^ = 0.03, *F*(5,71) = 0.44, *p* = 0.82). When using sleep CoV as the dependent variable, the full regression model was significant (R^2^ = 0.237, *F*(5,71) = 3.38, *p* = 0.001) and, the variation reduced with age (*β* = -0.01, *t*(71) = -2.08, *p* = 0.04) and increased with weight (*β* = 0.003, *t*(71) = 3.02, *p* = 0.004). Next, we performed the same analysis using the sleep duration and sleep CoV values derived from tappigraphy. For the sleep duration, the overall model was highly significant (R^2^ = 0.30, *F*(5,71) = 6.17, *p* = 8.39 × 10^−^). The higher the phone usage the shorter the duration (*β* = -0.0002, *t*(71) = -4.71, *p* = 1.2 × 10^−5^) and the larger the weight the shorter the duration (*β* = -0.023, *t*(71) = -2.09, *p* = 0.04). For sleep CoV, the overall model was significant (R^2^ = 0.23, *F*(5,71) = 4.25, *p* = 002). The higher the phone use the lower the variation (*β* = -1.7 × 10^−5^, *t*(71) = -3.19, *p* = 0.002) and females were less variable than males (female = 1, *β* = -0.09, *t*(71) = -3.03, *p* = 0.003).

### Smartphone usage in actigraphy derived ‘sleep’

Some of the observations described above were consistent with the idea that actigraphy can overestimate sleep. If this is the case, smartphone touches must be visible even during the putative sleep times determined by actigraphy. First, we quantified the probability of observing a smartphone touch during the actigraphy derived sleep. Users were found to be regularly interacting on the phone in the putative sleep period (Fig. 4). Next, we addressed the temporal distribution of the probability of observing smartphone touches in 3 min bins after the sleep onset and before the wake-up time. Interestingly, the probability of observing a touch remained significantly greater than 0 for ≈ 2 h after sleep onset and before wake-up time. Unsurprisingly, the body movements were observed through the night, albeit lower than at sleep onset or wake-up time.

**Figure 4.**
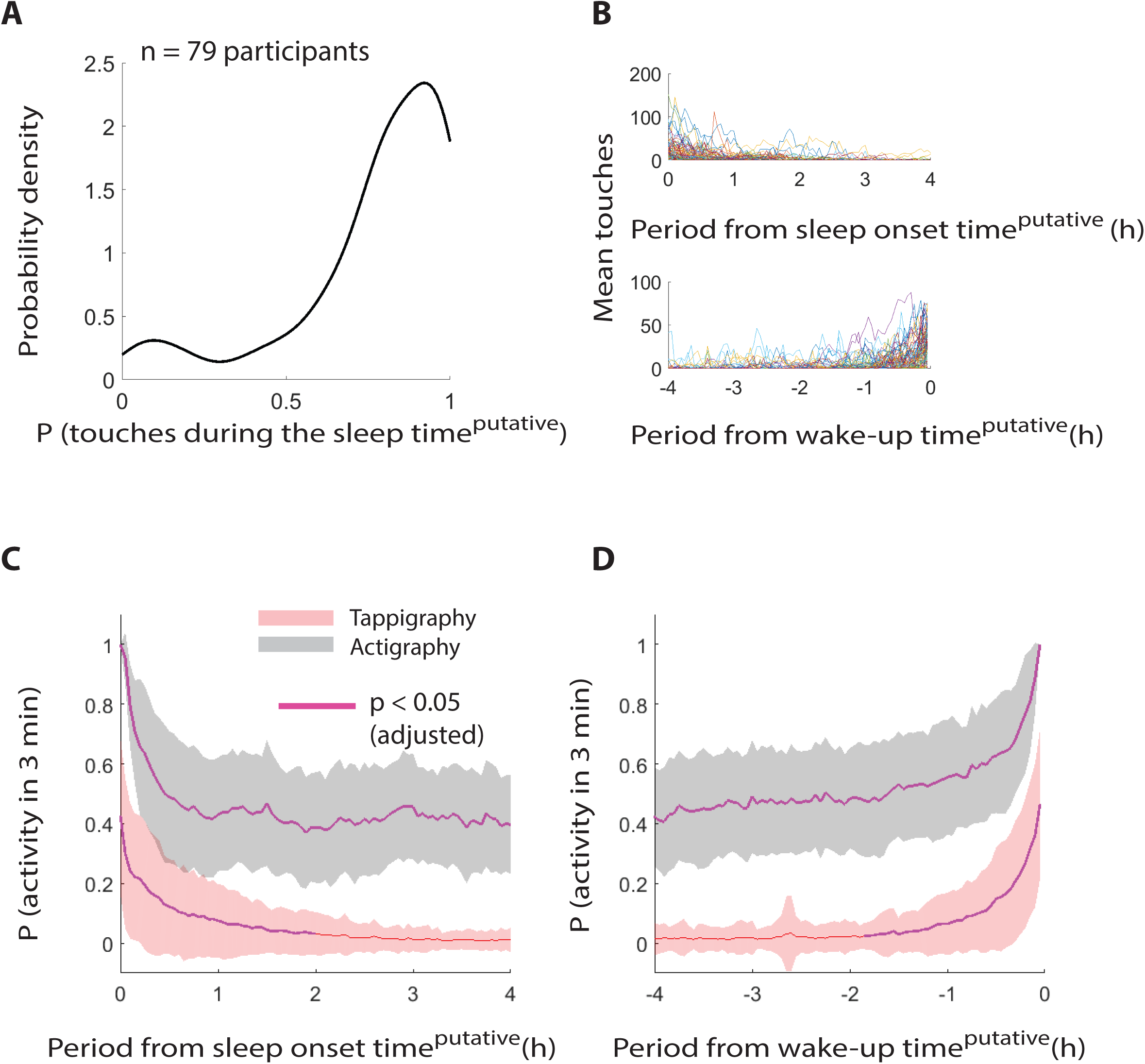
Prevalence of smartphone touches during actigraphy inferred sleep. (A) Kernel density plot of the probability of observing smartphone touches during ‘sleep’ in the sampled population – based on the percentage of nights with smartphone touches extracted from each participant. (B) The break-down of the number of touches generated after sleep-onset and before wake-up times with each participant represented using a different line. The probability of detecting smartphone touches and gross body movements in 3 min bins after sleep onset (C) and before wake-up time (D). The dark line represents the central tendency of the population and the shaded area represents the standard deviation. Note, according the central tendencies smartphone touches are observed with a probability of ~0.4 right after sleep onset and right before wake-up time. The statistical testing against a probability of 0 was corrected for multiple comparison correction.

## Discussion

This study revealed the state of physical activity during the cognitively engaging fine movements captured on the smartphone touchscreen. The smartphone touches consistently occupied the periods of low physical activity and this has substantial repercussions for sleep. On the one hand it offers a new opportunity to quantify sleep and on the other hand, it offers a highly quantitative insight into the extent of integration of modern digital behaviour with sleep behaviour, in particular with falling asleep and waking up. According to our results, smartphone touches recorded in the background can yield a reliable proxy measure of sleep and surprisingly, the digital interactions are as much a part of falling asleep as they are a part of waking up.

Deriving sleep-related measures from the day-to-day smartphone interactions may fundamentally improve health care. First, it offers an economical sleep screening tool that can be implemented in large populations with no user effort. Moreover, the method can be run in the background for prolonged durations (weeks to years) to trace the development of sleep. The touchscreen interactions remove the reliance on using gross movements to monitor sleep in the real world – and this can help quantify sleep in clinical populations where gross movements are severely attenuated. For instance, patients instructed to undergo bed-rest in the hospital or at home may not generate sufficient movements for actigraphy but may continue to engage in smartphone interactions^13^. Similarly, there are a plethora of clinical conditions – from obesity to arthritis – where fine movements are sufficiently preserved for smartphone interactions but the overall mobility is diminished^14,15^. Therefore, smartphone interactions may offer a new window into sleep in clinical populations where actigraphy or polysomnography cannot be easily implemented.

Firmly understanding the link between smartphone usage and physical activity is a key step for sleep research and medical conditions associated with physical inactivity or obesity^16^. For the former, there is no consensus on to what extent smartphone usage or digital media usage influences sleep but the conventional observations have vigorously employed self-reports which can be expected to provide a noisier, more time consuming and expensive understanding compared to the sensors used here^17,18^. We consistently find that users used their phones at rest. In this study, we focused on a measure of physical activity that is typically used in resolving the sleep-wake state in the popular Cole-Kripke algorithm^3^. This measure uses the accelerations recorded at the wrist as raw inputs. A well-known limitation of actigraphy is that while low acceleration values with the watch firmly attached on to the wrist is a reliable indicator of physical rest, the higher values may be contaminated for instance by a bumpy ride in a vehicle. Therefore, when considering our finding that there can be a second peak of smartphone usage when physically active must take this technical limitation into consideration.

We deployed a simplistic algorithm to determine the putative sleep onset and wake-up times based on the smartphone touches. This tappigraphy algorithm essentially combined two safe assumptions. First, the smartphone screen can be only touched when awake. Second, users follow a 24-hour sleep-wake cycle. The first assumption provided us with a list of smartphone usage gaps of which at least one contained sleep duration. The second assumption helped select the maximum gap which overlapped with the inactive phase – and this gap was identified as putative sleep. This simple approach resulted in sleep onset and wake-up times which were highly correlated with the times extracted from the standard actigraphy or sleep diary. Admittedly, there is scope for improvement – neither do we foresee a solution to how tappigraphy could detect day-time naps nor can we be sure it would accurately reflect sleep in subjects who cannot follow the 24 h cycle as in shift workers or in sleeping disorders such as insomnia. However, this initial version can be a powerful tool for quantifying sleep in individuals who follow diurnal behavioural patterns and addressing its utility in shift workers or sleeping disorders is a necessary next step.

Interestingly, the sleep durations were typically shorter when measured by using tappigraphy in comparison to actigraphy or sleep diaries. The overestimation of sleep by actigraphy is a well-recognized methodological issue and is typically explained by the delay between reduced motility and falling asleep^3,13^. Our findings suggest a more complex scenario in the sense that there is a highly active period – in terms of cognitively engaging smartphone behaviour – between the two sleep-related landmarks. A surprising finding was the prevalence of smartphone interactions surrounding the wake-up times, suggesting another source of sleep-overestimation in actigraphy. In sum, smartphones occupy the apparently quiescent periods between going to bed and falling asleep, and waking up and getting out of bed. These findings raise the crucial question of whether these periods were used differently in terms of cognitive activity prior to the introduction of smartphones in human behaviour. Regardless, due to the general consensus that a quiescent period before sleep is integral to initiating sleep, combining tappigraphy and actigraphy (or polysomnography) may offer highly relevant measures of sleep hygiene^19^.

We opportunistically used the demographic data assimilated on age, gender, height and weight to address how they related to sleep measures derived from tappigraphy and actigraphy. In terms of sleep duration, actigraphy based values did not relate to the demographic information. However, according to tappigraphy the higher body weight (height-adjusted) negatively correlated with sleep duration. This is in line with previous findings on obesity and sleep, and at the very least phone usage surrounding sleep may be indicative of body weight^14^. An interesting pattern of results emerged when we focused on the night-to-night variations in sleep using tappigraphy or actigraphy. Again, in line with previous observations on irregular sleep in obesity, actigraphy revealed that individuals with higher weight showed more irregularity in sleep. In tappigraphy, we found a striking gender difference, with females being more regular sleepers than males. The findings from such a small sample are by no means conclusive, but it does offer an interesting starting point in using these methods to address inter-individual differences and underscores that tappigraphy may be sensitive to distinct features compared to actigraphy.

Smartphones are truly integrated into modern human behaviour, including in our sleeping habits. Quantifying the extent of integration may not only yield a better understanding of behaviour in the real world but also yield new measures of sleep. The sleep measures introduced here do not rely on high smartphone usage as such but rather rely on the phenomenon that smartphones are used at rest and that they can be easily used even when in bed. The favourable consequence of this deep digital integration is that we can now develop highly scalable measures of sleep. Whether the integration is determinantal to sleep itself needs to be clarified.

**Author contributions**
AG and RH designed the study. AG analysed the data and acquired the data with the help of JB. AG drafted the manuscript with the help of RH.

## Acknowledgements

We thank the students in the Masters in Applied Psychology program (D. van Berkel, M. Dekker, E. van Duijn, K. van Eijgen, M. van Eijk, N. Grootscholten, K. Nieuwenhuizen, A. Tran, T. van Tuijl and M. van Weerwijk) for their help in data collection and recruitment, and the students in the Bachelors program (A.C. de Groot, A. Stam, I. Vedder and L. Klok) for their help in transcribing the sleep diaries. This work was supported by the intra-mural funding by the Unit of Cognitive Psychology at Leiden University, the Swiss National Science Foundation (320030_153387), the Clinical Research Priority Program (CRPP) Sleep and Health of the University of Zurich and the Hochschulmedizin Zurich (HMZ) Flagship grant SleepLoop. We thank Anna Kutschireiter for coining the term ‘Tappigraphy’.

## Notes

**Conflict of interest** Arko Ghosh is a co-founder of QuantActions Ltd, Lausanne, Switzerland. This company focuses on converting smartphone taps to mental health indicators. Software and data collection services from QuantActions was used to monitor smartphone activity.

